# Variable branching characteristics of peripheral taste neurons indicates differential convergence

**DOI:** 10.1101/2020.08.20.260059

**Authors:** Tao Huang, Lisa C. Ohman, Anna V. Clements, Zachary D. Whiddon, Robin F. Krimm

## Abstract

Taste neurons are functionally and molecularly diverse, but their morphological diversity remains completely unexplored. Using sparse cell genetic labeling, we provide the first reconstructions of peripheral taste neurons. The branching characteristics across 96 taste neurons show surprising diversity in their complexities. Individual neurons had 1 to 17 separate arbors entering between 1 to 7 taste buds, 18 of these neurons also innervated non-taste epithelia. Axon branching characteristics are similar in gustatory neurons from male and female mice. Cluster analysis separated the neurons into four groups according to branch complexity. The primary difference between clusters was the amount of the nerve fiber within the taste bud available to contact taste-transducing cells. Consistently, we found that the maximum number of taste-transducing cells capable of providing convergent input onto individual gustatory neurons varied with a range of 1-22 taste-transducing cells. Differences in branching characteristics across neurons indicate that some neurons likely receive input from a larger number of taste-transducing cells than other neurons (differential convergence). By dividing neurons into two groups based on the type of taste-transducing cell most contacted, we found that neurons contacting primarily sour transducing cells were more heavily branched than those contacting primarily sweet/bitter/umami transducing cells. This suggests that neuron morphologies may differ across functional taste quality. However, the considerable remaining variability within each group also suggests differential convergence within each functional taste quality. Each possibility has functional implications for the system.

**Significance statement:** Taste neurons are considered relay cells, communicating information from taste-transducing cells to the brain, without variation in morphology. By reconstructing peripheral taste neuron morphologies for the first time, we found that some peripheral gustatory neurons are simply branched, and can receive input from only a few taste-transducing cells. Other taste neurons are heavily branched, contacting many more taste-transducing cells than simply branched neurons. Based on the type of taste-transducing cell contacted, branching characteristics are predicted to differ across (and within) quality types (sweet/bitter/umami vs sour). Therefore, functional differences between neurons likely depends on the number of taste-transducing cells providing input and not just the type of cell providing input.

## Introduction

The chemical information from food is detected by cells in the taste bud (taste-transducing cells) and carried to the brain by sensory neurons. This chemical information includes taste qualities described as sweet, salty, sour, bitter, and umami (Contreras and Lundy 2000, Spector and Travers 2005, Yarmolinsky, Zuker et al. 2009, Carleton, Accolla et al. 2010, Ohla, Yoshida et al. 2019) as well as those that are not easily classified (Bachmanov, Tordoff et al. 1996, Tordoff 2001, Liu, Archer et al. 2016, Tordoff 2017, Lim and Pullicin 2019). Moreover, taste stimuli vary in intensity, and differences in stimulus intensity impacts quality coding (Ganchrow and Erickson 1970, Wu, Dvoryanchikov et al. 2015). Adding to their functional diversity, taste neurons respond to and are modulated by somatosensory stimuli (Lundy and Contreras 1997, Breza, Curtis et al. 2006, Yokota and Bradley 2016, Yokota and Bradley 2017). While numerous studies have focused on the functional diversity of this population of neurons, only recently has there been confirmation of their molecular diversity (Dvoryanchikov, Hernandez et al. 2017, Zhang, Jin et al. 2019). However, their morphological diversity remains completely unexplored.

Unlike the sensory neurons of skin (Maksimovic, Nakatani et al. 2014, Castillo, Diaz-Franulic et al. 2018), taste neurons do not participate in transduction (Chandrashekar, Hoon et al. 2006). Therefore, functional differences between gustatory neurons are thought to be determined entirely by the type of taste-transducing cell they innervate (Yarmolinsky, Zuker et al. 2009). Consistent with this hypothesis, retrograde tracing studies conclude that most taste neurons innervate only a single taste bud (Zaidi and Whitehead 2006). In addition, functional studies conclude that limited convergence is required for quality coding at mid-range stimulus concentrations (Yoshida, Yasumatsu et al. 2006). Despite the functional and molecular diversity of taste neurons, anatomical variation in these neurons was not expected, and was difficult to measure.

A definitive measure of morphological diversity necessitates full reconstruction of the peripheral axon of individual neurons. However, anatomical tracers do not travel the full length of peripheral taste axons. Genetic labeling provides a practical alternative (Wu, Williams et al. 2012, Bai, Lehnert et al. 2015, Kuehn, Meltzer et al. 2019); however, obstacles to this approach also exist. First, the number of taste neurons innervating the tongue is miniscule relative to the somatosensory innervation of lingual epithelium. Thus, if neurons from both populations are labeled, individual taste axons will be indistinguishable from somatosensory axons. As these two populations of neurons express many of the same factors, the majority of genetic labels used in other systems are impractical for reconstructing taste neurons (Wu, Williams et al. 2012). Lastly, the tongue is an unusually dense tissue requiring relatively thin sections for antibody labeling, which renders full neuron reconstruction unusually tedious.

It is important to overcome these practical limitations to fully reconstruct the axons of taste neurons for several reasons. Classification of morphological diversity can be combined with molecular expression data to correlate morphologies with distinct functions (Abraira and Ginty 2013). Knowing whether morphological differences are present dictates whether this direction of inquiry would be informative for the field of taste. Furthermore, the degree of branching likely reflects the degree of connectivity with taste-transducing cells (Chklovskii 2004), which would indicate that gustatory neuron responses are determined by both the type and number of transducing cells providing input.

In the present study, we used genetically directed sparse-cell labeling to fully reconstruct the complete peripheral morphologies of 96 neurons innervating taste buds in the tongue. Quantitative analyses revealed a surprising degree of diversity between taste neurons that, on average, innervate more taste buds than previously estimated (Zaidi and Whitehead 2006). We demonstrate that neurons with more complex axonal arbors contact a large number of taste-transducing cells, whereas those with simple endings contact only a few taste-transducing cells. We conclude that individual neurons likely receive differing amounts of convergent information from taste-transducing cells.

## MATERIALS AND METHODS

### Animals

*TrkB*^CreER^ mice (*Ntrk2*^*tm3*.*1(cre/ERT2)Ddg*^;, https://www.jax.org/ ISMR Cat# JAX:027214, RRID:IMSR_JAX:027214) were crossed with Cre-dependent alkaline phosphatase (AP) mice (https://www.jax.org/; IMSR Cat# JAX:009253, RRID:IMSR_JAX:009253) to obtain TrkB^CreER^:AP mice, or with Cre-dependent tdTomato mice ((Rutlin, Ho et al. 2014) https://www.jax.org/ RRID: IMSR_JAX:007914) to obtain *TrkB*^CreER^:tdTomato mice. In these mice, either the reporter gene AP or tdTomato is expressed following TrkB-driven Cre-mediated gene recombination. Similarly, Phox2b-Cre mice (MRRC_034613-UCD) were also bred with tdTomato reporter mice, such that all taste neurons were labeled with tdTomato..

Fungiform taste buds decrease in number postnatally (Liebl, Mbiene et al. 1999, Patel and Krimm 2012), such that varying the experimental age of examination could introduce variation in fiber morphology. To avoid this potential confounding variable, all fibers were reconstructed in mice at postnatal day 60–62. Although all animals were examined at the same age, 27 *TrkB*^CreER^:AP mice were injected with tamoxifen (0.5mg-1.0mg) at P40, axons were reconstructed from 17 of these 27 mice (8 male and 9 female), while 10 either had no labeled axons or too many labeled axons to reconstruct (more than 4). To select from all neurons expressing TrkB during development, 48 pregnant dams were injected with 4-hydroxytamoxifen at E15.5, 26 litters survived through birth, producing 21 mice (9 male and 12 female) that were of the correct genotype and had the correct number of axons to permit reconstruction (1-4), 115 mice either had 0 labeled axons or too many to accurately reconstruct (more than 4). To avoid selection bias, either none of the fibers were reconstructed or all of the fibers were reconstructed for each half tongue. In order to label one single tdTomato-positive axon in the tongue, 94 *TrkB*^CreER^:tdTomato mice were injected with tamoxifen (0.3mg-1mg), producing a total of 21 half tongues with one labeled axon. The remaining 167 tongue halves were determined to have either no labeled axons or too many (more than one) following serial sectioning of the tongue muscle. All animals were cared for in accordance with the guidelines set by the U.S. Public Health Service Policy on the Humane Care and Use of Laboratory Animals and the NIH Guide for the Care and Use of Laboratory Animals.

### Tamoxifen injections

Tamoxifen (T-5648, Sigma, St. Louis, MO) was dissolved in corn oil (C-8267, Sigma, St. Louis, MO) at 20 mg/ml by shaking and heating at 42°C and injected at P40 by intragastric gavage. For embryonic injections, 4-Hydroxytamoxifen (4-HOT) (H-7904, Sigma, St. Louis, MO), the active metabolite of tamoxifen, was injected intraperitoneally (i.p.) into pregnant dams at E15.5. 4-HOT was prepared and injected as previously reported (Wu, Williams et al. 2012). Briefly, 4-HOT was dissolved in ethanol at 20 mg/ml by shaking and incubating at 37°C for 15 min and then stored at -20°C. Before use, the stock solution was dissolved in sunflower seed oil to a final concentration of 10 mg/ml 4-HOT, and the ethanol was evaporated by centrifugation under vacuum.

### Alkaline phosphatase (AP) staining

Mouse tongues were dissected following perfusion with PBS containing 2% paraformaldehyde (PFA), 0.5% glutaraldehyde, and 2 mM MgCl_2_. The tongues were cut in half and postfixed for 2 h. After rinsing twice in PBS with 2 mM MgCl_2_, the tissues were transferred to PBS without MgCl_2_, and incubated at 65°C for 2 h to inactivate endogenous alkaline phosphatase (AP). Then, the tongues were frozen in O.C.T. (Sakura Finetek, USA) and stored at -80°C before serial sectioning at 180 µm. The staining was performed according to a previous publication with slight modification (Badea, Wang et al. 2003). Briefly, the tongue sections were washed in 0.1 M Tris-HCl (pH 7.5) followed by 0.1 M Tris-HCl (pH 9.5) before incubating overnight at room temperature in 0.1 M Tris-HCl (pH 9.5) containing nitroblue tetrazolium and 5-bromo-4-chloro-3-indolyl-phosphate at the recommended concentrations (Vector Laboratories). After washing in PBS and postfixing in 4% PFA, the stained tissues were dehydrated with an ethanol series, cleared and mounted with 2:1 benzyl benzoate/benzyl alcohol (Sigma); sections were sealed with DPX (VWR).

### Chorda tympani nerve transection

Mice were sedated with a 0.32-mg/kg intramuscular injection of medetomidine hydrochloride (Domitor) and anesthetized with 40 mg/kg ketamine hydrochloride (Ketaset). Mice were placed in a nontraumatic head holder to provide access to the nerve in the neck via a ventral approach (Guagliardo and Hill 2007). The chorda tympani nerve was located as it bifurcates from the lingual branch of the trigeminal nerve and was transected (portion was removed) without damaging the trigeminal nerve. The wound was sutured, and mice recovered on a water-circulating heating pad before being returned to their home cages. Atipamezole hydrochloride (2 mg/kg) was injected intramuscularly immediately after surgery to promote the reversal of anesthesia and thus reduce recovery time. Meloxicam was also administered orally through food pellets for 2 days after surgery; to achieve a dose of 1–2 mg/kg, 0.5 cc of a 0.5 mg/ml meloxicam solution was applied to each of two food pellets, which were then moistened with water and placed in the home cage. After 7 days, mice were euthanized and perfused with 4% PFA.

### Fluorescent anterograde nerve labeling

The procedures used to label the chorda tympani with fluorescent tracers were previously described (Sun, Dayal et al. 2015). Briefly, adult *TrkB*^CreER^:tdTomato mice, which had been given 4.0 mg tamoxifen via gavage for 3 weeks, were anesthetized and placed in the head holder as described above. A water-circulating heating pad was used to maintain body temperature. The chorda tympani nerves within the right tympanic bulla were cut near and peripheral to the geniculate ganglion, and crystals of 3 kDa fluorescein dextran (D3306; Invitrogen) were applied to the proximal cut end of the chorda tympani. A small amount of Kwik-Sil (World Precision Instruments, Inc., Sarasota, FL) was then placed over the cut ends of the nerves to prevent crystals from diffusing from the intended labeling site. Postsurgical treatment was the same as described above. After 24 h, mice were euthanized and perfused with 4% PFA. The geniculate ganglia were dissected and immediately mounted and imaged by confocal microscopy.

### Immunohistochemistry

*TrkB*^CreER^:tdTomato mice were sacrificed by avertin overdose (4 mg/kg) and perfused transcardially with 4% PFA. Dissected tissues were postfixed in 4% PFA for 2 h (for thin serial sections) or overnight (thick sections and whole mounts), rinsed with PBS, and transferred to 30% sucrose at 4°C overnight. A razor blade was used to remove the circumvallate papilla; the tongues were then carefully split down the midline with a razor blade under a dissection microscope. Tongues were frozen the next day in OCT and stored at -80°C before sectioning on a cryostat or processing for whole-mount staining. To determine if tongue halves are innervated by a single fiber, the muscle at the base of the midline was isolated using a razor blade, frozen in OCT, and serially sectioned (30 µm) on the cryostat. Serial sections were thaw mounted onto slides in order and then cover slipped with Fluoromount-G (SouthernBiotech, Birmingham, AL). Sections were examined under a fluorescence microscope to determine how many labeled fibers enter the tongue.

To visualize taste buds and their innervation in serial sagittal sections, each tongue half was sectioned sagittally from the midline to the lateral edge at 90 µm. Each section was processed in a separate well, so that the section order was maintained. Floating sections were blocked at 4°C in 0.1 M PB with 3% donkey serum and 0.5% Triton X-100 overnight, and then incubated for 5 days at 4°C with primary antibodies, rabbit anti-DsRed (1:500, RRID:AB_632496, Living Colors DsRed polyclonal; Takara Bio USA) and rat monoclonal anti-Troma1 (cytokeratin 8, 1:50, RRID:AB_531826; Developmental Studies Hybridoma Bank, Iowa City, IA). Sections were then rinsed four times for 15 min each, incubated for 2 days in secondary antibodies, Alexa 488 anti-rat IgG (1:500; Jackson ImmunoResearch Labs Cat# 712-545-153, RRID:AB_2340684) and Alexa 555 anti-rabbit IgG (1:500;Molecular Probes Cat# A-31572, RRID:AB_162543), rinsed another four times for 15 min each, and then mounted and coverslipped.

Whole-mount immunohistochemistry of the lingual epithelium was performed to visualize innervated taste buds. First, the underlying muscle and lamina propria were removed as described previously (Ohman-Gault, Huang et al. 2017). The isolated lingual epithelium was then washed for 15 min (3 times) in 0.1 M PB. Tissues were then incubated in blocking solution (3% donkey serum, 0.5% Triton X-100 in 0.1 M PB) at 4°C overnight and then incubated for 5 days at 4°C with primary antibodies (PLCβ2, 1:500, RRID:AB_2630573; Santa Cruz Biotechnology) in antibody solution (0.5% Triton X-100 in 0.1 M PB). Tissues were rinsed four times for 15 min each with 0.1 M PB, incubated with secondary antibodies (1:500, Alexa Fluor 488 AffiniPure, RRID:AB_2340619; Jackson ImmunoResearch), rinsed again (4 times for 15 min each with 0.1 M PB), and then incubated with 5% normal rabbit serum in antibody solution. Tissues were then rinsed and incubated with AffiniPure Fab fragment donkey anti-rabbit IgG (20 µg/mL, RRID:AB_2340587; Jackson ImmunoResearch) in antibody solution, rinsed, and incubated with Zenon Alexa Fluor 555 rabbit IgG labeling kit (according to the instructions for Zenon complex formation [Z25305; Invitrogen]) using anti-DsRed (1:500; RRID:AB_10013483; Living Colors DsRed polyclonal; Takara Bio USA). Tissues were rinsed, incubated for 5 days at 4°C with Car4 primary antibody (1:500, RRID:AB_10013483; R&D Systems), rinsed, and then incubated with secondary antibodies (1:500, Alexa Fluor 647 AffiniPure, RRID:AB_2340438). Tissues were then rinsed again, mounted with Fluoromount-G, and coverslipped (high precision, 0107242; Marienfeld).

### Confocal Imaging

Taste bud images were obtained using an Olympus Fluoview FV1000 confocal laser-scanning microscope with a 60× NA1.4 lens objective using a zoom of 3, Kalman 2. Image sizes were initially set at 1,024 × 1,024 pixels but were cropped to reduce scanning time and bleaching. Serial optical sections at intervals of 0.47 μm in the Z dimension were captured, which is the optimal size at 60x magnification for 3D reconstruction. All colors were imaged sequentially in separate channels to avoid bleed through. Image stacks were then deconvolved using AutoQuant X3 software (Media Cybernetics, Maryland) to reduce out-of-focus florescence and improve image quality.

### Experimental Design and Statistical Analyses

#### Image analysis

AP-stained taste axons were reconstructed under a microscope (using a 60× lens objective with a 2-mm working distance and numerical aperture of 1.0) using Neurolucida software (MBF Bioscience). The axons were reconstructed from where the axons enter the tongue to all terminal branch ends. Axons were only reconstructed from a tongue half when all the axons in the tongue half could be reconstructed. Typically, only half tongues with fewer than 4 labeled axons could be reconstructed. Thus, axons from half tongues with more than 4 labeled axons were not reconstructed. Individual arbors innervating whole immunostained taste buds were also reconstructed from confocal image stacks using Neurolucida software. Reconstructions were then analyzed using Neurolucida Explorer (MBF Bioscience).

Imaris software version 6.4.2 (Bitplane) was used to generate the 3D reconstructions and to measure proximity of nerve arbors to labeled taste cells. The colocalization function in Imaris was not used, because it contains many user-selected options that might contribute experimenter bias. More importantly, it was unclear how much colocalization there would be between labels in two separate cell types: the labeled taste bud cells and nerve arbors. Also, the physical relationship between any two florescent markers in a sample is influenced by tissue processing, intensity and wavelength of labels, the location of the labeled protein in the cells, deconvolution, orientation of the tissue, etc. Thus, the physical distance between nerve arbors and labeled taste bud cells (proximity) was measured with a distance transformation function in Imaris, which was previously described (Valm, Cohen et al. 2017). This algorithm permits identification of an object and then its distance from any/all other objects in a defined 3D space. This was accomplished using automated thresholding, which identified the surface of labeled objects (cells and nerve arbors) and then determined the distance between them in voxel increments. Thresholds were automatically generated with no input from the operator to limit bias. Because the sampling was at roughly twice the resolution of the microscope, distances of two voxels or less are equivalent to the colocalization artifacts that occur when two objects of different colors are sufficiently close (Corson and Erisir 2013, Stratford, Larson et al. 2017). However, the proximity analysis enables the distance between any two cells to be measured. To display the relationship between a single arbor and labeled taste bud cell, individual taste-transducing cells and arbors were segmented in Imaris, and the fluorescent channel was only duplicated within the selected region. Segmentation was completed after analysis for illustrative purposes only.

#### Data analysis

Sixteen different anatomical measures were used in the k-means cluster analysis to group neurons on the basis of similarity: age of the animal at the time of tamoxifen injection, number of taste buds innervated, total length of the peripheral axon, location in the tongue of innervated taste buds (back, middle, tip), number of branch points (nodes) below the epithelium, number of branch points (nodes) in the taste bud, total number of branch ends, number of terminal branch ends per taste bud, mean terminal branch length, combined terminal branch length, combined arbor length, number of arbors, number of widened endings, number of non-taste endings, distance from the base of the tongue to the first branch point, and highest branch order (Fig. 1). Time of tamoxifen injection and sex of the animals were compared for each measure using either a Mann-Whitney U test or t-test depending on whether the measure was normally distributed (tested with Shapiro-Wilk test). A Bonferroni correction factor for the number of tests (16) required a p=0.003 to statistical significance. Cluster analysis was performed using MATLAB (https://www.mathworks.com/help/stats/k-means-clustering.html?s_tid=srchtitle). We tested 2 to 6 clusters and selected the 4-cluster model because it was the highest silhouette value with no negative values (second highest overall). Characteristics were compared across clusters by first testing whether the measure was normally distributed across all 4 clusters using the Shapiro-Wilk test. If the measure was normally distributed, then differences were determined using a one-way analysis of variance with a Bonferroni’s *post hoc* analysis. If they were not normally distributed differences across multiple groups were compared using a Kruskal-Wallis test, while two groups were compared using a Mann-Whitney test. Multiple comparisons were avoided by comparing a limited number of factors relevant to the study. Because most of our measures were not normally distributed, Spearman correlations were used to analyze the relationship between variables. The alpha level was set at *p*=0.05, and actual *p* values are reported. However, when more than one comparison was made for a measure, a Bonferroni’s correction factor was used to adjust the alpha level.

**Figure 1.**
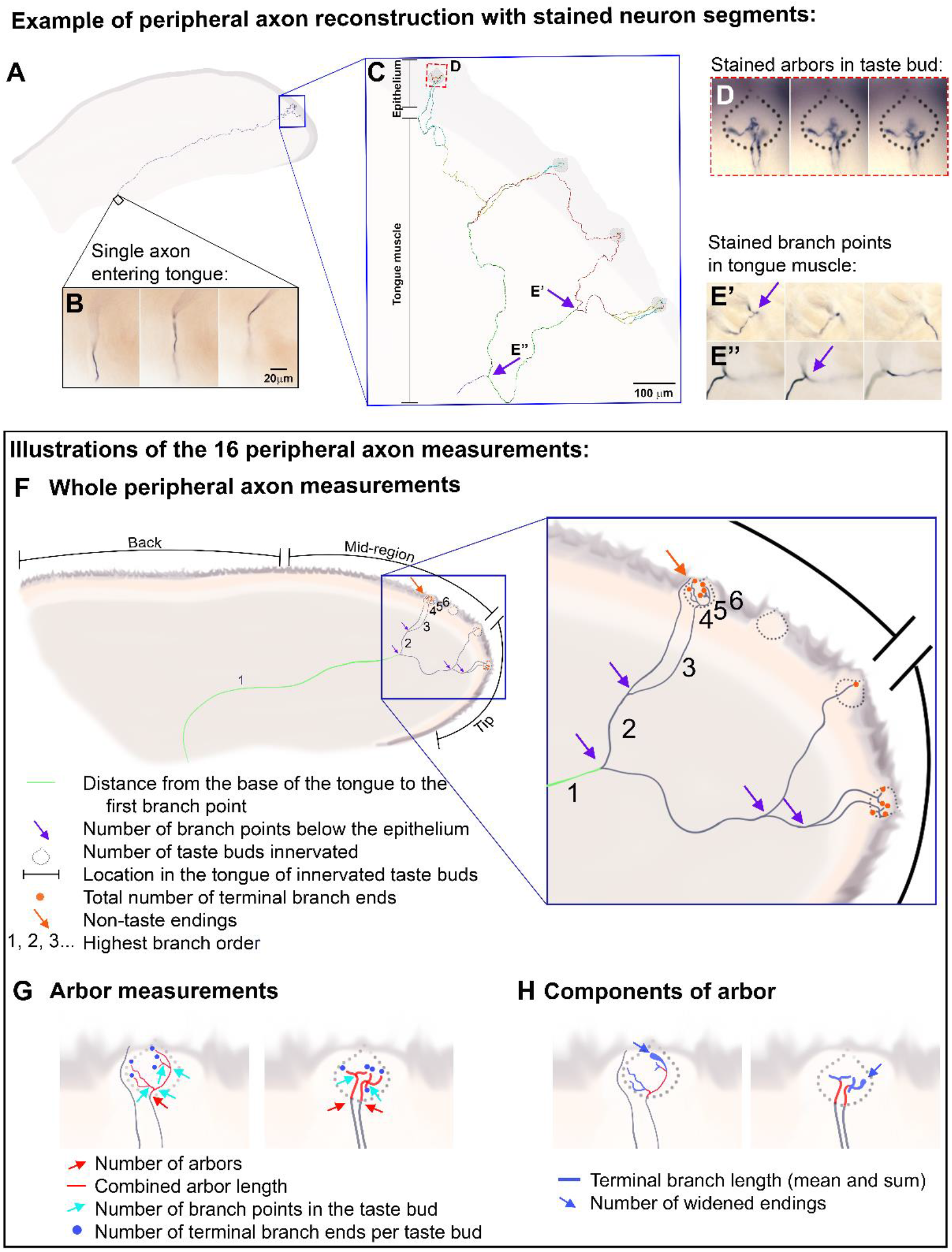
Genetically directed sparse-cell labeling, reconstruction, and quantification of individual taste axons. (A-E) A single AP-stained axon from a *TrkB*^CreER^:AP mouse injected with tamoxifen at postnatal day 40. (A) A reconstruction of the taste axon starting from where it enters the tongue (B) to the arbors. It is superimposed on an outline of a tongue section. (B) Images of the AP-stained axon are shown at 3 different focal depths (left to right) in this 180-µm section. (C) An enlarged view of the same axon with each branch segment presented in a different color. This axon innervates 4 taste buds (one of which is shown in D). Boxes illustrate the locations of branch points (E’-E’’), which are shown at 3 different focal depths. The scale bar in B = 20µm and applies to D and E; the scale bar in C = 100µm. (F) illustrates 7 of the features quantified for the entire axon (in the legend), using the mean features across all 96 axons to create this depiction. In addition, to the depicted measure the total length of the axon was measured (green and gray in F summed). The location in the tongue was assigned a number (tip=1, mid-region=2, back=3) was assigned, since the depicted hypothetical axon has arbors innervating both the tip and mid-region, the number 1.5 would be assigned. (G-H) Illustrate 7 additional anatomical characteristics measured in the taste buds.

## Results

### Genetically directed sparse labeling of taste neurons

Taste neurons are functionally and molecularly diverse, but it is unclear whether there is any morphological diversity within this population of neurons. To evaluate morphological diversity, we used an inducible genetic technique that primarily labels taste neurons, and not the more abundant somatosensory neurons that innervate the tongue epithelium. The neurotrophin receptor TrkB is expressed in most neurons within the geniculate (taste) ganglion (Huang and Krimm 2010, Tang, Rios-Pilier et al. 2017, Rios-Pilier and Krimm 2019), but very few TrkB-positive trigeminal neurons innervate the tongue (Wu, Arris et al. 2018). Therefore, we used *TrkB*^CreER^:alkaline phosphatase (AP) mice to reconstruct single taste nerve axons innervating the tongue (Rutlin, Ho et al. 2014). AP has been used to reconstruct single axonal arbors of somatosensory neurons (Badea, Williams et al. 2012, Wu, Williams et al. 2012), and it has the advantage of being stained with adequate intensity to be visualized in thick tissues, allowing the stained axons to be more readily reconstructed than with fluorescent labeling.

We injected 27 postnatal day 40 *TrkB*^CreER^:AP mice with low doses of tamoxifen to label a small number of individual neurons by adulthood (postnatal day 60) and reconstructed the peripheral axons from where the chorda-lingual nerve entered the tongue to where they terminated at the epithelial surface (Fig. 1A-E). The number of neurons labeled was variable and not strictly dose dependent. When more than four axons were labeled, they crossed paths, making it impossible to follow individual axons. To avoid the inherent bias of choosing the easiest axons to reconstruct when multiple axons were labeled, no reconstructions were completed from half tongues with more than four labeled axons. However, at the same doses of tamoxifen some tongues did not contain even a single labeled axon, while others had more than 4 labeled neurons. We reconstructed a total of 41 quantifiable neurons from 17 animals of the 27 injected with tamoxifen.

Since TrkB expression decreases in the geniculate ganglion after embryonic day 15.5 (E15.5) (Rios-Pilier and Krimm 2019), we were concerned that injections at postnatal day 40 did not include the full population of taste neurons. Therefore, we injected *TrkB*^CreER^:tdTomato mice with a high dose of tamoxifen (4mg/day for two weeks) to capture as many TrkB-expressing neurons as possible and then labeled the taste neurons projecting to the tongue with a fluorescent tracer (Sun, Dayal et al. 2015). We found that 73% (n=2, 71%, 75%) of tongue-innervating geniculate ganglion neurons were tdTomato-positive. Thus, the majority of TrkB-expressing taste neurons are labeled when adult *TrkB*^CreER^:tdTomato mice are injected with tamoxifen (Rios-Pilier and Krimm 2019). Nevertheless, to circumvent the reduction of TrkB during development, we injected 58 pregnant dams (at E15.5) with 0.5, 0.75, or 1.0 mg 4-Hydroxytamoxifen. A total of 55 axons were reconstructed from 21 of the 115 mice injected at E15.5. A total of 94 mice had either no labeled axons or more than 4 labeled axons, and so no axons were reconstructed from these mice.

All reconstructed axons were from tongues of adult mice (postnatal day 60–62) regardless of the age at injection. The age at time of tamoxifen injection P40 (n=41) and E15.5 (n=55) had no significant effect on 16 measured morphological characteristics (Fig. 1F-H,Table 1a.) There were no differences in most of the morphological characteristics of axons reconstructed from male (*n*=45) and female (*n*=51) mice (Table 1b.) The length of the axon from where it enters the tongue to the first branch point was slightly longer in males (9.32mm) than females (7.7mm, U =721, *p*=0.002), perhaps because the tongues were longer in males. Since differences based on age of tamoxifen injection and sex of the animal were minimal, we combined these data to yield a total of 96 fully reconstructed peripheral taste nerve axons (available to the research community at neuromorpho.org/dableFiles/tao_krimm/Supplementary/Tao_Krimm.zip). Across these neurons, we observed that individual axons had branch points in the tongue muscle (Fig.1E), lamina propria, and epithelium. Branch points below the epithelium dictated the number of separate arbors (Fig.1F), the portion of the axon innervating taste buds (range, 1–17 per neuron), whereas branch points inside the taste bud dictated the variation in complexity of these different arbors.

**Table 1:** Statistical Table. Significance level was set initially at p ≤ 0.05, but then adjusted for the number of tests using a Bonferroni correction (Sig. level).

|  | Measure | Data structure | Comparison | Type of test | Result | p-value | Sig. level |
| --- | --- | --- | --- | --- | --- | --- | --- |
| a. | number of taste buds | non-normal | day of injection | Mann-Whitney | U = 1011 | p=0.36 | p≤0.003 |
| | total axon length | normal | day of injection | t-test | $t_{(94)}=0.69$ | p=0.49 | p≤0.003 |
|  | location in the tongue | non-normal | day of injection | Mann-Whitney | U=962 | p=0.203 | p≤0.003 |
|  | #of branch points below the epithelium | non-normal | day of injection | Mann-Whitney | U=798 | p=0.013 | p≤0.003 |
|  | number of branch points in the taste bud | non-normal | day of injection | Mann-Whitney | U=962 | p=0.2 | p≤0.003 |
|  | average number of branch ends/bud | non-normal | day of injection | Mann-Whitney | U = 750 | p=0.005 | p≤0.003 |
|  | total number of branch ends | non-normal | day of injection | Mann-Whitney | U=1006 | p=0.35 | p≤0.003 |
|  | average arbor length per taste bud | non-normal | day of injection | Mann-Whitney | U=1008 | p=.309 | p≤0.003 |
|  | combined terminal branch length | non-normal | day of injection | Mann-Whitney | U=987 | p=0.28 | p≤0.003 |
| | mean terminal branch length per bud | normal | day of injection | t-test | $t_{(94)}=-0.28$ | p=0.78 | p≤0.003 |
|  | number of arbors | non-normal | day of injection | Mann-Whitney | U=968 | p=0.22 | p≤0.003 |
|  | combined arbor length | non-normal | day of injection | Mann-Whitney | U = 960 | p=0.2 | p≤0.003 |
|  | number of widened endings | non-normal | day of injection | Mann-Whitney | U=899 | p=0.82 | p≤0.003 |
|  | length of axon from entry to first branch point | non-normal | day of injection | Mann-Whitney | U=812 | p=0.02 | p≤0.003 |
|  | highest branch order | non-normal | day of injection | Mann-Whitney | U=1089 | p=0.738 | p≤0.003 |
|  | number of non-taste endings | non-normal | day of injection | Mann-Whitney | U=1127 | p=0.944 | p≤0.003 |
| b | number of taste buds | non-normal | male vs. female | Mann-Whitney | U = 885 | p=0.05 | p≤0.003 |
| | total axon length | normal | male vs. female | t-test | $t_{(94)}=0.42$ | p=0.69 | p≤0.003 |
|  | location in the tongue | non-normal | male vs. female | Mann-Whitney | U=1025 | p=0.35 | p≤0.003 |
|  | #of branch points below the epithelium | non-normal | male vs. female | Mann-Whitney | U=792 | p=0.007 | p≤0.003 |
|  | number of branch points in the taste bud | non-normal | male vs. female | Mann-Whitney | U=1044 | p=0.45 | p≤0.003 |
|  | average number of branch ends/bud | non-normal | male vs. female | Mann-Whitney | U=1111 | p=0.74 | p≤0.003 |
|  | total number of branch ends | non-normal | male vs. female | Mann-Whitney | U=883 | p=0.052 | p≤0.003 |
|  | average arbor length per taste bud | non-normal | male vs. female | Mann-Whitney | U=1008 | p=.309 | p≤0.003 |
|  | combined terminal branch length | non-normal | male vs. female | Mann-Whitney | U = 980 | p=0.22 | p≤0.003 |
| | mean terminal branch length per bud | normal | male vs. female | t-test | $t_{(94)}=-0.88$ | p=0.38 | p≤0.003 |
|  | number of arbors | non-normal | male vs. female | Mann-Whitney | U = 812 | p=0.013 | p≤0.003 |
|  | combined arbor length | non-normal | male vs. female | Mann-Whitney | U = 953 | p=0.15 | p≤0.003 |
|  | number of widened endings | non-normal | male vs. female | Mann-Whitney | U = 1131 | p=0.91 | p≤0.003 |
|  | length of axon from entry to first branch point | non-normal | male vs. female | Mann-Whitney | U=721 | p=0.002 | p≤0.003 |
|  | highest branch order | non-normal | male vs. female | Mann-Whitney | U=820 | p=0.015 | p≤0.003 |
|  | number of non-taste endings | non-normal | male vs. female | Mann-Whitney | U=1073 | p=0.91 | p≤0.003 |
| Fig.4F | combined arbor length | normal | clusters | ANOVA | $F_{(3,92)}=368$ | p=0.001 | p≤0.05 |
| Fig.5C | axon length vs number of taste buds innervated | non-normal | cluster 2 | Spearman correlation | $\rho=0.68$ | p=0.0000002 | p≤0.05 |
| Fig.5E | # of arbors vs. mean length per arbor | non-normal | Cluster 2 | Spearman correlation | $\rho=-0.61$ | p<0.001 | p≤0.05 |
|  | branch points in muscle and lamina propria | non-normal | cluster 1 vs cluster 2 | Mann-Whitney | U=254 | p=0.001 | p≤0.008 |
| Fig.6C | number of taste buds innervated | non-normal | cluster 1 vs cluster 2 | Mann-Whitney | U=296 | p=0.001 | p≤0.008 |
| Fig.6D | axon length | normal | cluster 1 vs cluster 2 | t-test | $t_{(75)}=4.72$ | p=0.001 | p≤0.008 |
|  | terminal branch ends per arbor | non-normal | cluster 1 vs cluster 2 | Mann-Whitney | U=742 | p=0.89 | p≤0.008 |
|  | terminal arbor length | non-normal | cluster 1 vs cluster 2 | Mann-Whitney | U=575 | p=0.07 | p≤0.008 |
| Fig.6F | number of taste buds vs combined arbor length | non-normal | cluster 1 vs cluster 2 | Spearman correlation | $\rho=-0.94$ | p=0.0000002 | p≤0.008 |
| Fig.7 | taste bud number | normal | cluster 2 vs cluster 3/4 | t-test | $t_{(52)}=-1.445$ | p=0.154 | p≤0.016 |
| Fig.7C | number of terminal arbors | non-normal | cluster 2 vs cluster 3/4 | Mann-Whitney | U=108 | p=0.001 | p≤0.016 |
| Fig.7D | terminal arbors/taste bud | non-normal | cluster 2 vs cluster 3/4 | Mann-Whitney | U=173 | p=0.006 | p≤0.016 |
| | mean arbor length | non-normal | clusters | Kruskal-Wallis | $H_{(2,93)}=7.18$ | $p=0.006$ | $p\leq 0.05$ |
| | number of arbors vs combined arbor length | non-normal | all axons | Spearman correlation | $\rho=0.75$ | $p=0.0000002$ | $p\leq 0.05$ |
| | arbor length | non-normal | arbors contacting 1 vs more than 1 | Mann-Whitney | $U=1185$ | $p=0.0001$ | $p\leq 0.025$ |
| | branch end number | non-normal | arbors contacting 1 vs more than 1 | Mann-Whitney | $U=1522$ | $p=0.0012$ | $p\leq 0.025$ |
| Fig.9D | number of taste-transducing cells contacted | non-normal | clusters | Kruskal-Wallis | $H_{(2,20)}=17.16$ | $p=0.0002$ | $p\leq 0.05$ |
| Fig.9E | patterns of innervation | non-normal | clusters 1, 2, and 3 | Chi-square | $\chi^2=6.63$ | $p=0.036$ | $p\leq 0.05$ |
| Fig.9F | arbor length | normal | axons contact more Car4+ vs PLCβ2+ cells | t-test | $t_{(19)}=2.59$ | $p=0.018$ | $p\leq 0.025$ |
| | branch ends | non-normal | axons contact more Car4+ vs PLCβ2+ cells | Mann-Whitney | $U=19$ | $p=0.025$ | $p\leq 0.025$ |

The large number of branch points in the musculature and lamina propria was surprising, because in mice most taste neurons are thought to innervate only a single taste bud. Because these earlier studies assumed that gustatory neurons innervate adjacent taste buds, we examined heavily branched neurons to determine if they always innervated adjacent papillae. Many neurons innervate immediately adjacent papillae. However, we also found some taste neurons innervating multiple taste buds that were not located in the adjacent fungiform papillae in the tongue (Fig. 2A). Approximately 10% of the neurons had these wide anatomic receptive fields. Unfortunately, because sections were thick and tissue other than nerve fibers were unstained, it was not possible to determine if all of them had uninnervated papillae between innervated papillae. Also, most neurons also tended to innervate each taste bud with only a single arbor (Fig. 2B), which tended to increase their receptive field sizes. For example, a neuron with 4 arbors innervating different taste buds would have a larger receptive field that a neuron with 4 arbors innervating a single taste bud.

**Figure 2.**
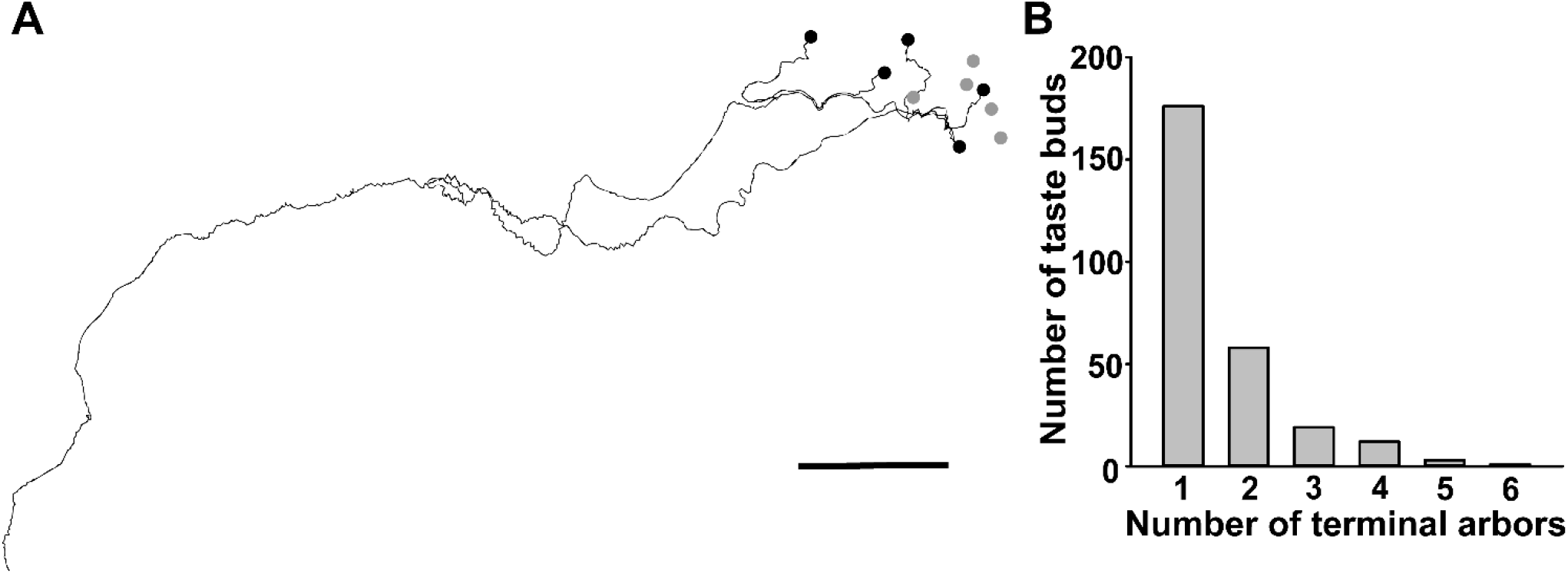
Anatomic receptive fields tend to be distributed over the lingual epithelial surface. (A) A sagittal view of a reconstructed axon that innervated 5 taste buds, with a total of 6 arbors (only one of these taste buds received more than one arbor). The fungiform papillae containing the innervated taste buds are marked by black dots. Other fungiform papillae containing taste buds not innervated by this neuron (marked by gray dots) and at least two are located between innervated papillae. Approximately 10% of axons have these extensive receptive fields. Scale bar = 1.0mm (B) A histogram illustrating how many arbors innervate each taste bud from any single neuron. The 96 reconstructed axons innervate a total of 279 taste buds, and most taste buds only receive a single arbor from each taste neuron.

In the 38 tongues from which the 96 TrkB-positive axons innervating taste buds were reconstructed, we also observed 4 labeled axons that did not innervate taste buds. To determine if all labeled axons innervating the front of the tongue were from the primary taste nerve (the chorda tympani nerve), we injected *TrkB*^CreER^:tdTomato mice (*n*=2) with a high dose of tamoxifen to label many TrkB-positive neurons and then transected the chorda tympani nerve. The axons remaining in the tongue after degeneration of the chorda tympani nerve are from another non-geniculate (taste) ganglion source (Guagliardo and Hill 2007). Although the number of TrkB-positive axons entering the tongue on the transected side was reduced compared to the contralateral side, some TrkB-positive axons remained in the tongue (Fig. 3A,B). These TrkB-positive axons innervating the tongue were possibly from the trigeminal ganglion (Wu, Arris et al. 2018). To determine if these non-chorda tympani TrkB-positive axons innervate taste buds, we quantified TrkB-positive innervation in the taste buds following unilateral chorda tympani nerve transaction on the intact side (Fig. 3C,D) compared to the transected side (Fig. 3E,F). We found that 100% of taste buds were innervated by tdTomato-labeled nerve axons on the intact side of the tongue, whereas only 5% of the taste buds were innervated by tdTomato-labeled nerve axons on the transected side. These innervated taste buds (four in two mice) were located at the midline ventral tongue tip where the midline is less clear, such that the innervation was likely from the contralateral chorda tympani nerve. We conclude that most TrkB-positive axons innervating the tongue are from the chorda tympani and that axons from other sources do not typically innervate taste buds. Therefore, axons that did not innervate taste buds were not reconstructed or included in the data set, as their nerve of origin was uncertain. So unfortunately, non-taste neurons within the chorda tympani (Dvoryanchikov, Hernandez et al. 2017, Yokota and Bradley 2017) nerve were probably not reconstructed.

**Figure 3.**
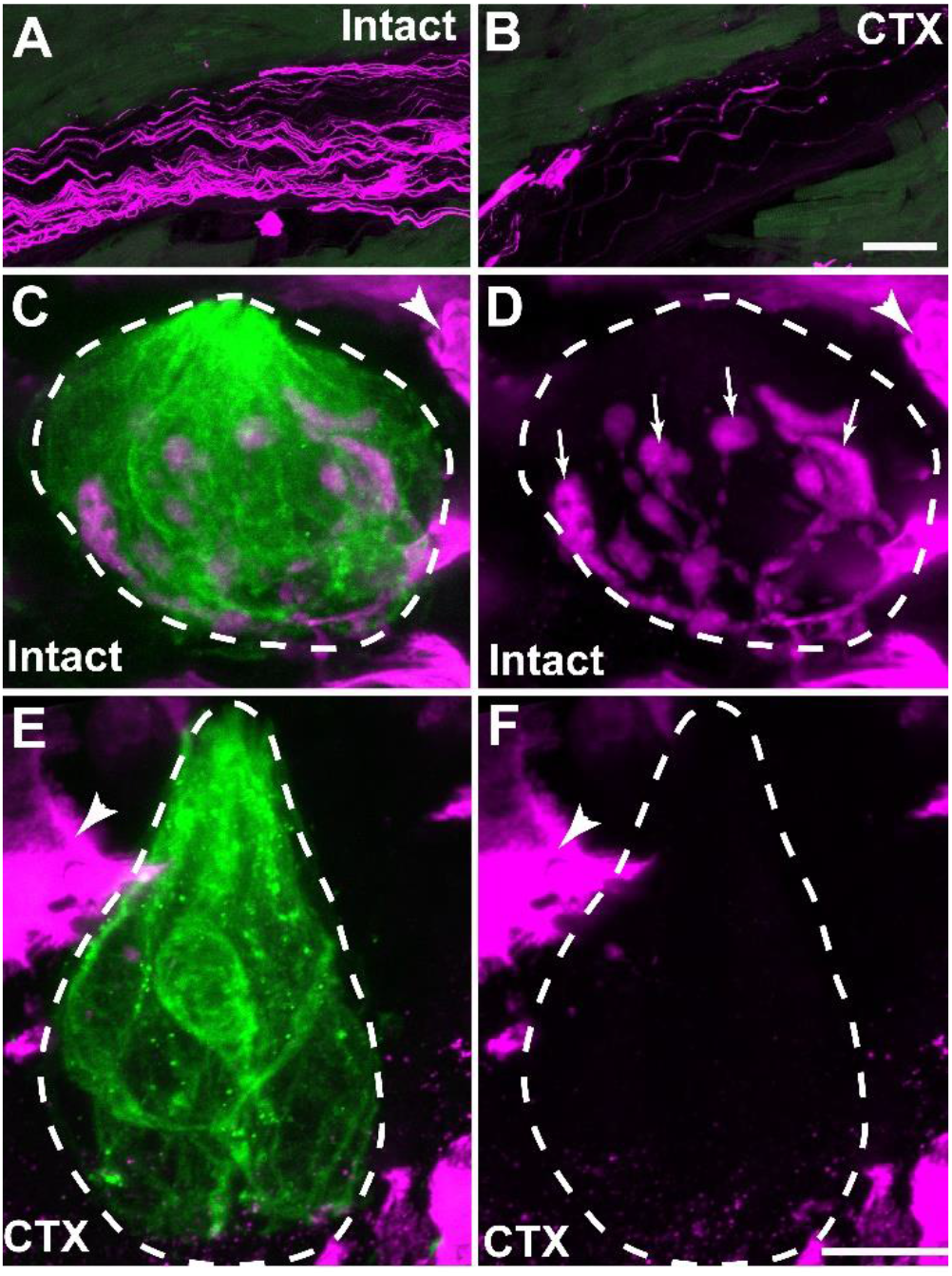
Most, but not all, TrkB-positive taste axons projecting to the tongue innervate taste buds. (A) On the un-transected side of the tongue (intact), abundant TrkB-positive axons (magenta) were observed in chorda-lingual nerve bundles. (B) On the chorda tympani transected side of the tongue (CTX) from the same mouse, substantially fewer TrkB-positive axons were observed in nerve bundles. (C-D) TrkB-positive axon nerve arbors can be seen in this example taste bud from the side of the tongue with the intact nerve. The taste bud was identified by cytokeratin-8 (green) which is enclosed with dashed line, arrows in D identify labeled arbors penetrating the taste bud shown in C. Because the truncated form of the TrkB receptor is expressed by keratinocytes, many keratinocytes outside the taste bud are also labeled with tdTomato because of high dose of tamoxifen used to label most taste neurons (arrowheads, C-F). (E,F) A taste bud from the CTX side of the tongue lacked labeled arbors inside the taste bud which was defined by cytokeratin 8 in E and outlined in F. Scale bar in B =50µm and also applies to A; scale bar in F = 10µm and also applies to C, D, and E.

### Individual axons of taste neurons vary in branching characteristics

Somatosensory neurons of the dorsal root ganglion can be classified as types by their distinct morphological endings in skin (Wu, Williams et al. 2012), this was not the case with taste neurons. However, we did observe that some taste axons branched very little, while others displayed extensive branching. To organize these neurons into descriptive groups (clusters), we measured 16 separate morphological characteristics (see Methods, Fig. 1F-H) and performed a k-means clustering analysis (Fig. 4). Silhouette values were used to estimate the optimal number of clusters. A four-cluster model provided the second highest mean silhouette value (Fig. 4A, 0.6944) while having no negative values, an indication of miss-assignment. We plotted histograms of all measured characteristics to determine which characteristics had the least amount of overlap between the clusters. The three characteristics allowing greatest separation between clusters are shown (Fig. 4B-D). These four clusters represented a gradual increase in branch complexity (measured morphological characteristics related to branching), from the simplest in cluster 1 to the most complex in cluster 4 (Fig. 4E). Just under half of the axons constituted cluster 1 (N=42), most of the remaining neurons in cluster 2 (N=38) and the fewest in the more complex cluster 3 (N=13) and cluster 4 (N=6). Combined arbor length in the taste bud was the only measured trait with no overlap between clusters and provides the clearest separation of the clusters (*F*_(3,92)_ = 368,*p*=0.001; Fig. 4F). This measure corresponds to the amount of the nerve fiber (axon) available to contact taste-transducing cells. This finding implies that number and not just type of taste-transducing cells providing input to taste neurons varies across the taste neuron population.

**Figure 4.**
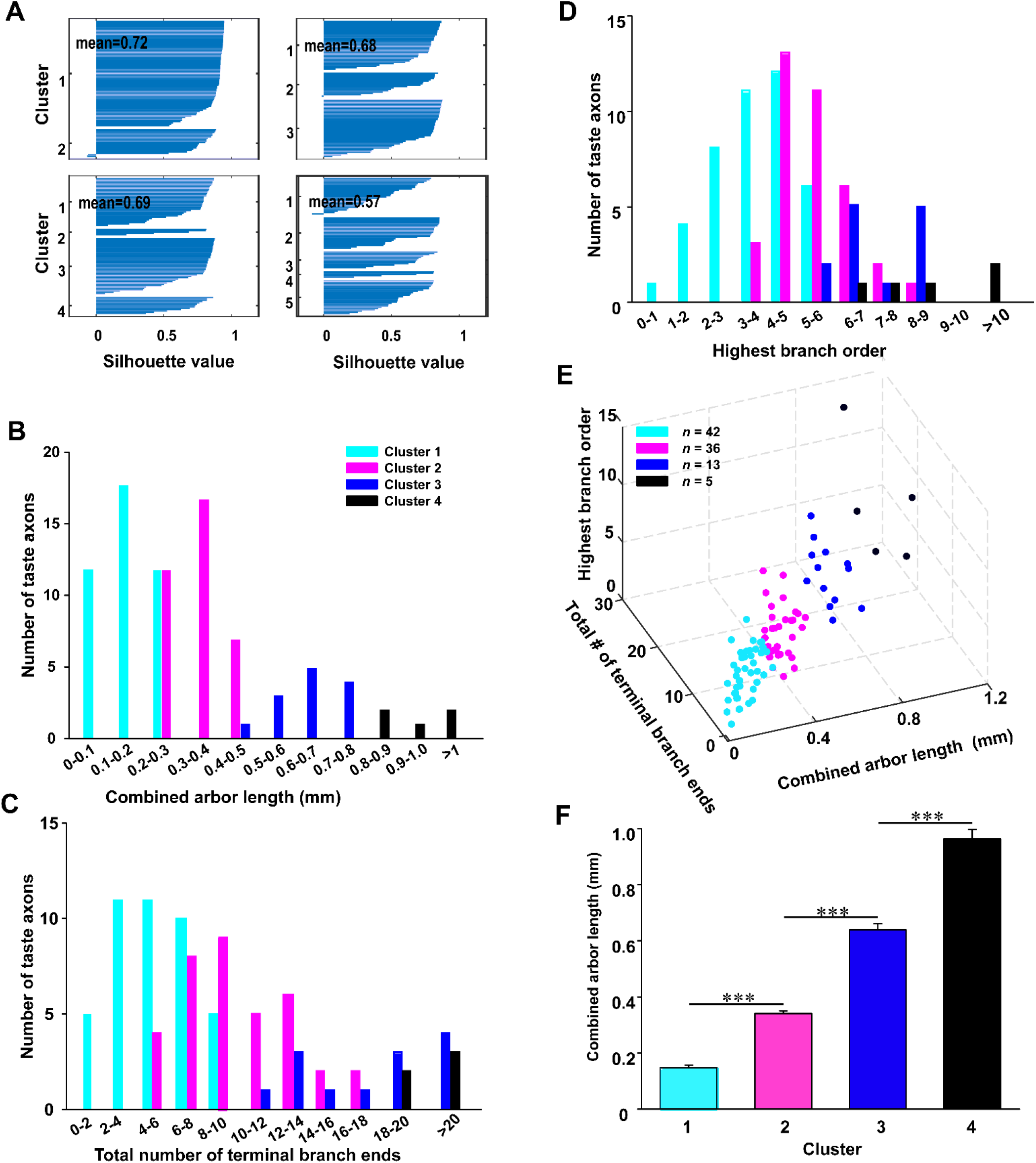
Taste axons were divided into four categories using a k-means cluster analysis. (A) Silhouette values are graphed for each neuron based on the similarity of each neuron to its own cluster (higher values = greater similarity). Four plots illustrate the number of clusters varying from 2 to 5. The highest silhouette value (0.72) was obtained assuming there were only two clusters, but the two-cluster model showed outliers. which also were observed in the cluster 3 and the cluster 5 models. A four cluster model had the second highest mean silhouette value (0.69) with no outliers. (B-D) The distributions of combined arbor length, total number of terminal branch ends, highest branch order. These are the measures showing the greatest separation between clusters. (E) 3D graph showing the relationship of these three characteristics for individual neurons. (F) Clusters are defined by combined arbor length, which is the amount of the axon available to form a connection with a taste-transducing cell.\*\*\**p* ≤ 0.001

### Nearly half (44%) of taste axons have simple peripheral morphologies

Representative examples of the simple morphologies of two cluster 1 neurons are shown in Figure 5A and B, which innervated one and two taste buds, respectively. Arbors (the portion of the axon inside the taste bud) of cluster 1 neurons typically had two to three (Fig. 5AB) branch points within the taste bud. Many arbors branched within the taste bud, with only a single branch point and two branch ends (Fig.5B first taste bud). Most of the axons in cluster 1 neurons innervated one or two taste buds (81%; Fig. 5C), with only one to two arbors (Fig. 5D). The more taste buds that are innervated the longer the axon (ρ=0.68, *p*=0.0000002, Fig. 5C). A smaller proportion (19%) of the cluster 1 axons innervated three or more taste buds (Fig. 5C) with four or more separate arbors (Fig. 5D). However, these arbors were unusually short (Fig. 5E). Although cluster 1 neurons are morphologically simple, all but one branched at least once (Fig. 5F). All regions of the lingual epithelium (Fig. 1F) were innervated by cluster 1 neurons. Specifically, eight neurons innervated taste buds located at the tongue tip, twelve neurons spanned between the tip and the mid-region, eleven neurons innervated the tongue mid-region only, two neurons innervated the back of the tongue, and three neurons spanned between the mid-region and the back of the tongue.

**Figure 5.**
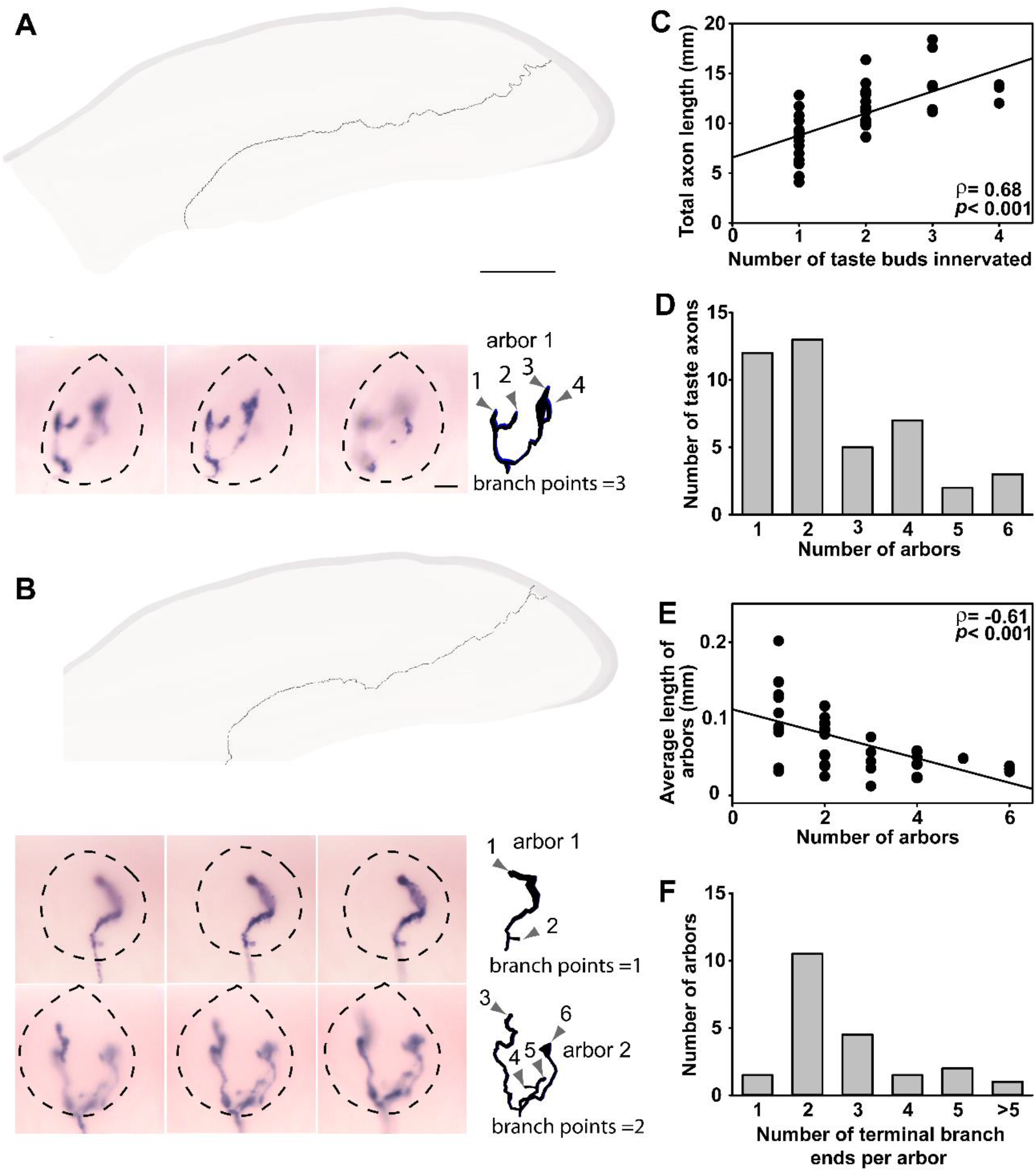
Two examples of axons from cluster 1 neurons. (A, B) The reconstructed axons were superimposed on an outline of one sagittal tongue section to illustrate the location of the axon in the tongue. The images below each reconstructed axon show the AP staining of arbors within the taste bud at three different focal depths. Reconstructions of the arbors within the taste bud are shown at the same magnification to the right of the images. (A) This 10.3-mm-long axon innervated a single taste bud with one arbor, three branch points inside the taste bud, and four terminal branch ends (gray arrowheads). (B) The second 14.0-mm-long axon innervated two taste buds with one arbor each. One branch point outside the taste bud and three inside the taste bud, results in six total terminal branch ends (gray arrowheads). (C) Axons from cluster 1 taste neurons innervate 1–4 taste buds. The number of taste buds that are innervated correlates with the total length of the axon. (D) The axons of cluster 1 neurons typically have 1 to 2 arbors penetrating taste buds, but could have as many as 6 arbors. (E) Axons of cluster 1 neurons with the most arbors also had on average the shortest arbors. (F) The distribution of the number of terminal branch endings per arbor. Most axons from cluster 1 neurons had arbors with 2 terminal branch ends. Scale bars: 1 mm for whole axon tracings in A and B; 10 µm for taste bud images and reconstructions within taste buds in A and B.

### Complex taste axons innervate more taste buds

Reconstructions of cluster 2 neurons illustrate an axon innervating three taste buds (Fig. 6A) and a second axon innervating seven taste buds (Fig. 6B). The neuron illustrated in Fig. 6A has widened endings (blue arrow), which were present in most taste nerve arbors; a feature that did not differ across clusters. Although the function of these endings is not known, flattened contacts (Romanov, Lasher et al. 2018, Yang, Dzowo et al. 2019) with taste bud cells or retraction bulbs are possibilities (Bernstein and Lichtman 1999, Zaidi, Cicchini et al. 2016). Compared to those in cluster 1, axons of cluster 2 neurons contained more branch points in the tongue muscle and lamina propria (median=1 (N=42) vs 4 (N=36), U=254, *p*=0.001) and innervated more taste buds (Fig. 6C, U=296, *p*=0.001). Thus, the entire axon innervating the tongue was longer on average for cluster 2 neurons than for cluster 1 neurons (t_(75)_=4.72, *p*=0.001, Fig. 6D). Similar to cluster 1, most of the axons of cluster 2 neurons had a single arbor innervating each taste bud (Fig. 6A and B). Cluster 1 and cluster 2 neurons did not differ in the number of terminal branch ends per arbor (median=2 (N=42) vs 2 (N=36), U=742, *p*=0.89) or arbor length (median=20.8 (N=42) vs 24.6 (N=36) µm, U=575, *p*=0.07). Therefore, the longer combined arbor length of cluster 2 axons is primarily due to an increase in the number of taste buds innervated (Fig. 6C,E). There were cluster 2 neurons that innervated only 1 or 2 taste buds; however, these neurons had longer arbors (*ρ*=-0.-94, p=0.0000002, Fig. 6F). Like cluster 1, the axons of cluster 2 neurons innervated all regions of the lingual epithelium, but they were more likely to innervate multiple tongue regions. Specifically, axons branching to innervate both the tongue tip and the mid-region (16 axons) were the most common, with fewer axons exclusively innervating the tongue tip (9) or mid-region (4). In addition, five cluster 2 neurons innervated the back of the tongue, and one innervated both the back and the mid-region.

**Figure 6.**
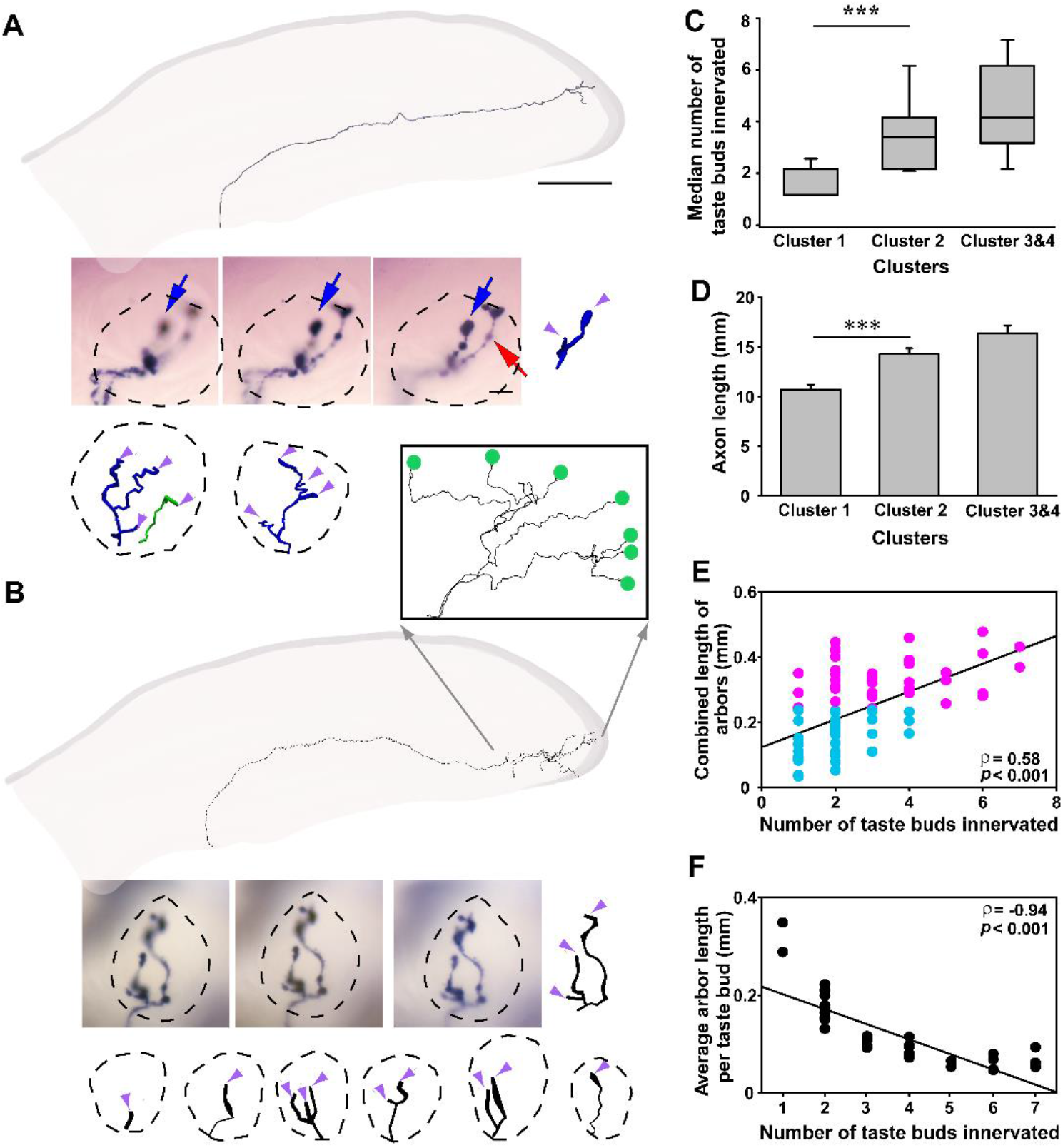
Two examples of axons from cluster 2 neurons. The reconstructed axons were superimposed on an outline of one sagittal tongue section to illustrate the location of the axon in the tongue. The images below each reconstructed axon show the AP staining of arbors within the taste bud at three different focal depths. Reconstructions of the arbors within the taste bud are shown at the same magnification to the right of the images. (A) The 13.0 mm axon of this cluster 2 neuron had 3 branch points below the epithelium producing 4 arbors that innervated 3 taste buds (second arbor in same taste bud is green). Inside the taste bud some of the arbors had additional branch points (5) to produce 10 terminal branch ends (purple arrowheads). The imaged taste bud has a single arbor (blue arrow) from the reconstructed axon plus one arbor from another labeled neuron (red arrow). Beneath the images are reconstructions of 3 additional arbors innervating two additional taste buds. (B) The 14.7 mm axon from a second neuron had 6 branch points below the epithelia, producing 7 arbors, each of which innervated a separate taste bud (reconstructions for 6 additional arbors are shown beneath the imaged arbor). These arbors not only innervated the dorsal and ventral tip, but also the lateral edge of the tongue. An inset shows the dorsal view of this axon (green dots indicate location of taste buds). Some of these arbors also branched in the taste bud (6 branch points) resulting in a total of 13 terminal branches (purple arrowheads). (C) Plots of medians ± interquartile ranges (IQR, gray boxes) and ± maximum/minimum (whiskers) illustrate that cluster 2 neurons innervated significantly more taste buds than cluster 1 neurons. (D) Similarly, the mean (± SEM) total length of axons from cluster 2 neurons was significantly greater than the axons from cluster 1 neurons, but not different from neurons in clusters 3 and 4. (E) As the number of taste buds a neuron innervated increased, the combined total length of the arbors also increased (cyan=cluster 1, magenta=cluster 2). The small number of cluster 2 neurons that innervated only a few taste buds (1-3), have longer arbors than cluster 1 neurons. (F) The average length of the arbors per taste bud for cluster 2 neurons decreased as more taste buds are innervated. Black line indicates linear fit. \*\*\**p*=0.001. Scale bars: 1 mm for whole nerve tracings in A and B; 10 µm for taste bud images and arbor reconstructions within taste buds in A and B.

### The most complex peripheral taste axons innervate taste buds with multiple arbors

Representative examples of axons from cluster 3 (Fig. 7A) and cluster 4 (Fig. 7B) neurons illustrate their greater complexity. Cluster 3/4 neurons innervate a similar number of taste buds as those in cluster 2 (Fig. 6C, t_(35,17)_ =-1.445, *p*=0.154), but with significantly more arbors (Fig. 7C, *p*=0.001). For example, seven taste buds were innervated by either 9 (Fig. 7A) or 10 (Fig. 7B) arbors. Similar to these examples, clusters 3 and 4 combined had more arbors per taste bud than cluster 2 neurons (Fig. 7D, U=173, *p*=0.006). Since the mean length of each arbor did not differ among the clusters (H_(2,93)_ =7.18, *p*=0.066), the combined length of the arbors inside the taste bud for an axon is primarily determined by number of arbors (ρ=0.75, *p*=0.0000002, Fig. 7E). The only difference between neurons in clusters 3 and 4 is the combined lengths of the arbors (Fig. 4F). Also, like axons of cluster 2 neurons, axons of cluster 3 and 4 neurons tended to innervate multiple tongue regions (9 out of 15).

**Figure 7.**
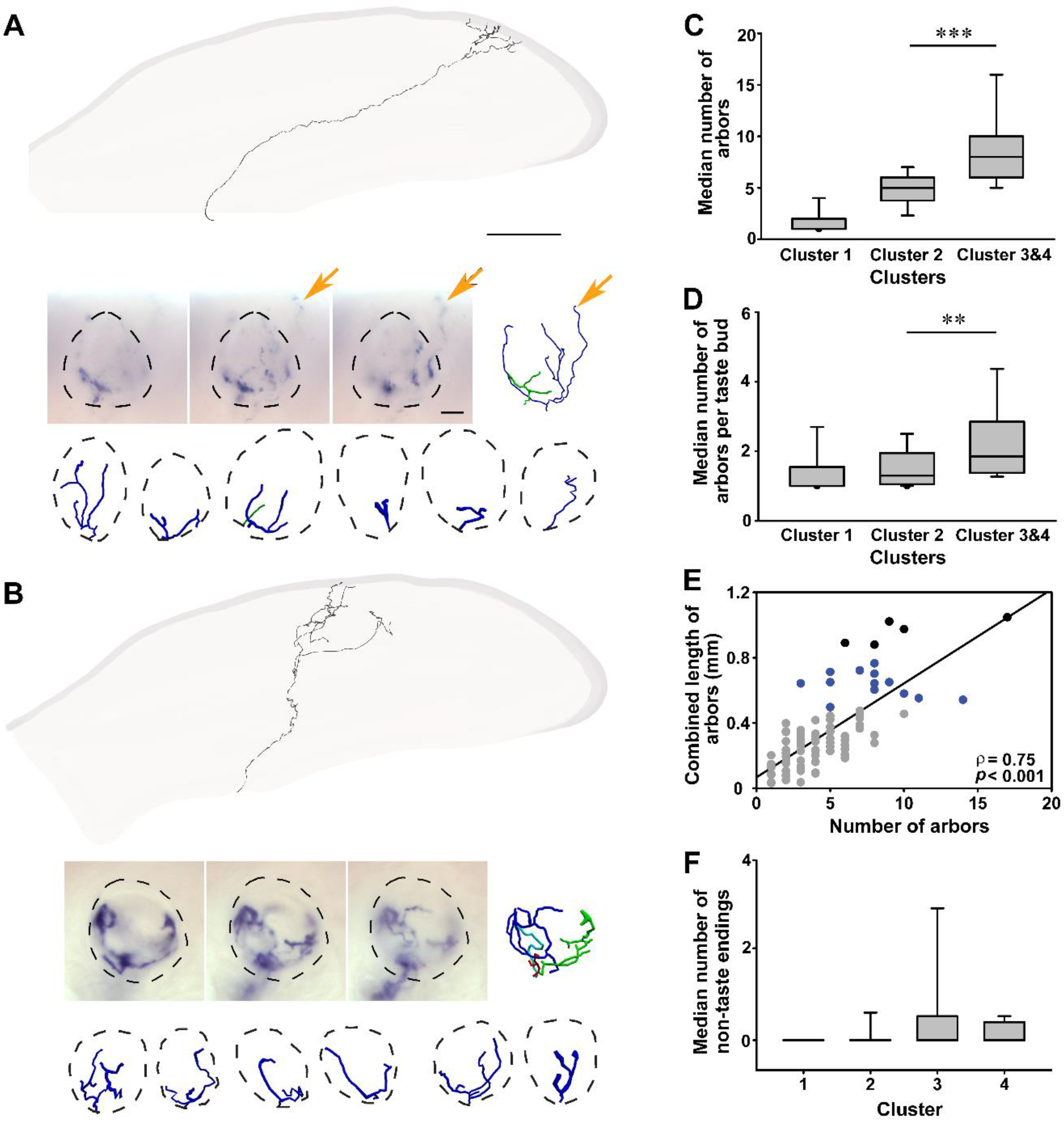
An example of an axon from a cluster 3 neuron and a cluster 4 neuron. Images below each axon reconstruction show the AP staining for one taste bud innervated by this axon and the reconstructions of all the arbors for this neuron. (A) The representative axon of a cluster 3 neuron had a total length of 18.6 mm, innervated 7 taste buds with 9 arbors, and had 28 terminal branches within the taste buds. Beneath the images are all the reconstructions of 7 additional arbors innervating 6 additional taste buds. For the two taste buds with two arbors the second arbor is shown in green. This axon also had an arbor innervating the epithelium outside of the taste bud (orange arrow, non-taste ending). (B) The axon of a cluster 4 neuron had a total length of 20.8 mm, innervated 7 taste buds, and had 10 arbors with total of 40 terminal branches. For this example, the imaged taste bud had 4 arbors (dark blue, green, red, cyan). Beneath the images are all the reconstructions of 6 additional arbors innervating 6 additional taste buds. (C-D) Plotted are medians ± interquartile ranges (IQR (gray boxes) and ± maximum/minimum (whiskers)). (C) The axons of cluster 3 and 4 neurons have more arbors than those in clusters 1 and 2. (D) This increase in arbor number is due to an increase in the average number of terminal arbors innervating each taste bud. (E) Across all neurons, the best indicator of the combined arbor length inside the taste bud is the number of arbors (blue=cluster 3, black=cluster 4, gray = other clusters). (F) A subset of neurons in each cluster had axons terminating outside of taste buds (medians and IQR, ± maximum/minimum). Scale bars: 1 mm for whole nerve tracings in A and B; 10 µm for taste bud images and reconstructions within taste buds in A and B.

All clusters had a few axons with branches ending outside of taste buds (Fig. 7A,F). Neurons with these “non-taste” endings typically have one or more arbors that penetrate the epithelium, with a free nerve ending in a region of the tongue not occupied by a taste bud. In some cases, these are entirely different arbors from those innervating a taste bud, but they could also branch from an arbor that innervates the taste bud. For example, the arbor of the first taste bud shown in Fig. 7A (dark blue) has a single branch arising from a branch point at the taste bud base (orange arrow) and extending into the lingual epithelium. A total of 18 neurons had these non-taste endings, which were most often in fungiform papillae (13 neurons) but also in filiform papillae (3 neurons) or both (2 neurons).

In summary, 44% of the peripheral axons of taste neurons had fairly simple branching characteristics (cluster 1), while the remaining 56% showed more complex branching. There were no differences in the average lengths or number of terminal branch ends of individual arbors across the 4 clusters, indicating that combined arbor length is determined primarily by the number of arbors and not their size. Axons of cluster 2 neurons increased arbor number by innervating more taste buds than cluster 1 neurons. Whereas axons of cluster 3 and 4 neurons increased arbor number by having more arbors per taste bud.

### Each arbor contacts a limited number of taste-transducing cells

Because clusters were best separated by the amount of axon available to contact taste-transducing cells, we next sought to examine how many taste-transducing cells are contacted by a single neuron. While it is not possible to observe connections at the light level, the largest number of taste-transducing cells that are sufficiently close to a neuron to form a connection can be determined. To determine the closest proximity between taste-transducing cells and nerve fibers, we labeled all taste nerve arbors (Phox2b-Cre:tdTomato mice) and many taste-transducing cells using well-established markers for cells transducing sweet, bitter, and umami-type stimuli [(anti-phospholipase C beta-2 (PLCβ2) (Zhao, Zhang et al. 2003, Clapp, Yang et al. 2004)] and those transducing sour-stimuli [(anti-carbonic anhydrase 4 (Car4) (Chandrashekar, Yarmolinsky et al. 2009)]. We then analyzed three-dimensional (3D) image stacks to identify the taste-transducing cells and arbors that were closest to each other (Fig. 8A). We found that arbors tended to be either within one voxel (∼110nm, below the resolution of the light microscope and frequently seen as overlapping (referred to henceforth as contacts) or separated by more than 1µm (Fig. 8A,B), so this became our criteria for a contact.

**Figure 8.**
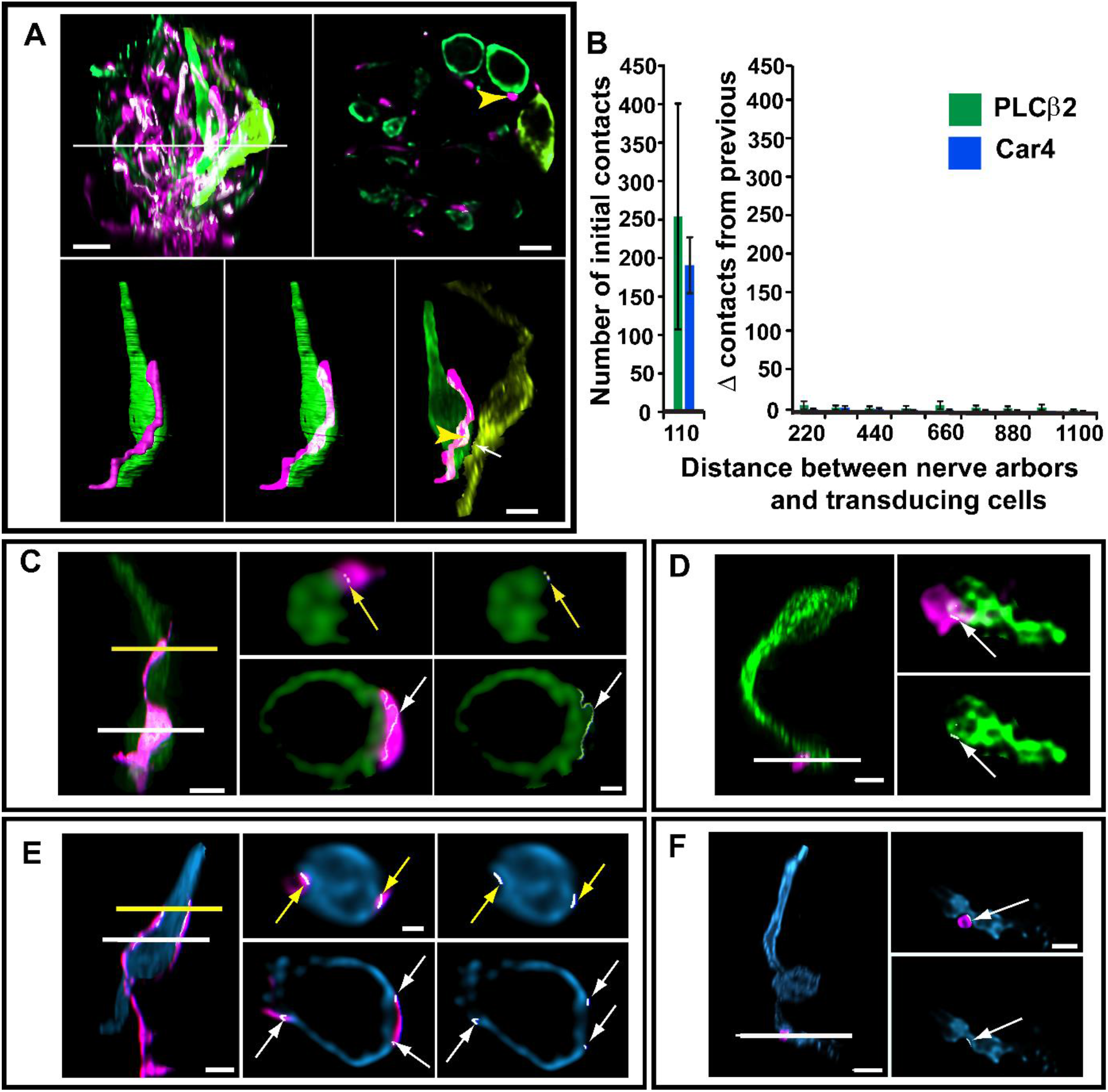
Determining the number of taste-transducing cells contacting individual arbors. (A) A taste bud with all arbors from taste neurons labeled magenta (tdTomato) and taste bud cells expressing PLCβ2-labeled green (sweet-, bitter-, and umami-transducing cells). The distance between nerve arbors and taste-transducing cells was measured incrementally in 1.5 voxels (110 nm, essentially overlapping). These regions were pseudo-colored white. A single section through the taste bud indicated by the white line is shown to the right of the taste bud. A single magenta arbor (yellow arrowhead) contacts one PLCβ2-positive taste bud cell (green). The next closest PLCβ2-positive cell to the same arbor was pseudo-colored yellow-green is 1.2µm away (white arrow). (A, bottom) Segmenting the cells and arbors to remove florescent label outside the segmented area permits the relationship between individual arbors and taste transducing cells to be viewed. Scale bar = 5µm. (B) When all innervation to the taste bud is labeled (using Phox2b-Cre:tdTomato), there are numerous locations where nerve arbors are within a single voxel (110 nm) of a taste-transducing cell (mean ± standard error for N=3 taste buds). However, very few additional cells are contacted as we increase the minimum distance (in nanometers), making 110 nm a reasonable criteria for a contact. (C-F) There are variations in the relationship between taste bud cells and nerve arbors. (C, E) Some arbors extend along a nerve cell for a long distance, others contact cells for shorter distances (D, F). Arbors tend to contact PLCβ2-labeled cells (green) at widened regions of the arbor (C, D white arrows). Arbors tend to contact Car4-positive cells (blue) at sites of indentation of the nerve fiber into the cell (E, F, arrows). The fluorescence for each fluorophore was optimized for brightness contrast. For panels C-F complete arbors and contacted taste-transducing cells were segmented and the fluorescence outside the reconstruction removed, as illustrated in panel A. The reconstruction was removed and the fluorescence inside the reconstruction is shown in each panel. Scale bars in C-F are 5µm in whole cell images, and 2µm in cross-sections.

To determine how many taste-transducing cells are contacted by individual taste arbors, we reconstructed 151 individual arbors in fungiform taste buds. These arbors were labeled in *TrkB*^CreER^:tdTomato mice injected with 1.5-2.0mg of tamoxifen, which tends to label 1 to 2 arbors in approximately half of the fungiform taste buds. Arbors contacting either PLCβ2-labeled (Fig. 8C,D) or Car4-labeled taste bud cells (Fig. 8E,F) sometimes followed the labeled taste bud cell a considerable distance (12-25µm; Fig. 8C,E), while others followed for much shorter distances (less than 12µm, Fig. 8D,F). Frequently, arbors contact the same cell at multiple different locations. Because it is unclear how many of these contacts actually represent a connection, we did not quantify contacts, but quantified the number of cells contacted. Thus, an arbor contacts one cell regardless of the distance travelled along the cell or the number of distinct instances in which the nerve arbor or taste receptor cell come within 200nm. Using this analysis, we found that each arbor contacted 1.87 taste-transducing cells on average. This number is only slightly more than the average of 1.6 Type III cells that synapse with individual circumvallate arbors (Kinnamon, Sherman et al. 1988).

We found that 48% of the arbors only contacted one taste-transducing cell, suggesting that these arbors either form a connection with a single taste-transducing cell or do not form a connection at all. A few of the arbors contacted no labeled taste bud cells (7%) and could not receive input from either a Car4-positive or PLCβ2-positive taste bud cell. Within the 45% that contacted multiple cells, 17% only contacted Car4- or only PLCβ2-positive cells such that 28% of the total population of arbors contacted both PLCβ2- and Car4-labeled taste bud cells. In addition to functional connections, an arbor likely contacts a taste-transducing cell when assessing molecular compatibility (Lee, Macpherson et al. 2017) to form a new connection or when a branch passes in proximity to a receptor cell. Because there is no published evidence that an individual arbor can receive functional input from more than one transducing cell type, we hypothesized that arbors contacting more than one cell type might be larger, and perhaps in a state of greater plasticity (process of connecting to a new cell). Consistent with this possibility, arbors contacting more than one cell type are longer (median=89 (N=106) vs 57 (N=39) µm, U=1185, *p*=0.0001) and more heavily branched (median 5 (N=106) vs 3 branch ends (N=39), U=1522, *p*=0.0012).

### Complex neurons contact more taste-transducing cells than simply branched neurons

Next, we sought to examine the degree to which more heavily branched neurons have an opportunity to connect with larger numbers of taste-transducing cells. To this end, we sought a dose of tamoxifen that would result in labeling single axons. Since these neurons typically do not cross the midline, each half of the tongue can be evaluated independently. Individual axons were identified from where the chorda tympani-lingual nerve enters the tongue in a single nerve bundle and followed through multiple serial sections to determine the number of labeled axons (Fig. 9A). We determined that one injection of 0.6 mg results in one single labeled axon roughly 25% of the time. Three examples of cluster 1 neurons are illustrated (Fig. 9B). The first neuron innervated one taste bud and contacted two Car4-positive cells. The second neuron innervated two taste buds and contacted 4 PLCβ2-positive cells, and the third neuron contacts one PLCβ2-postive cell. A single cluster 2 neuron is also illustrated (Fig. 9C), which contacted 10 labeled taste cells across 5 taste buds. Although this axon contacts multiple taste cell types, the contacts were heavily biased toward Car4-positive cells (7 Car4 and 3 PLCβ2).

**Figure 9.**
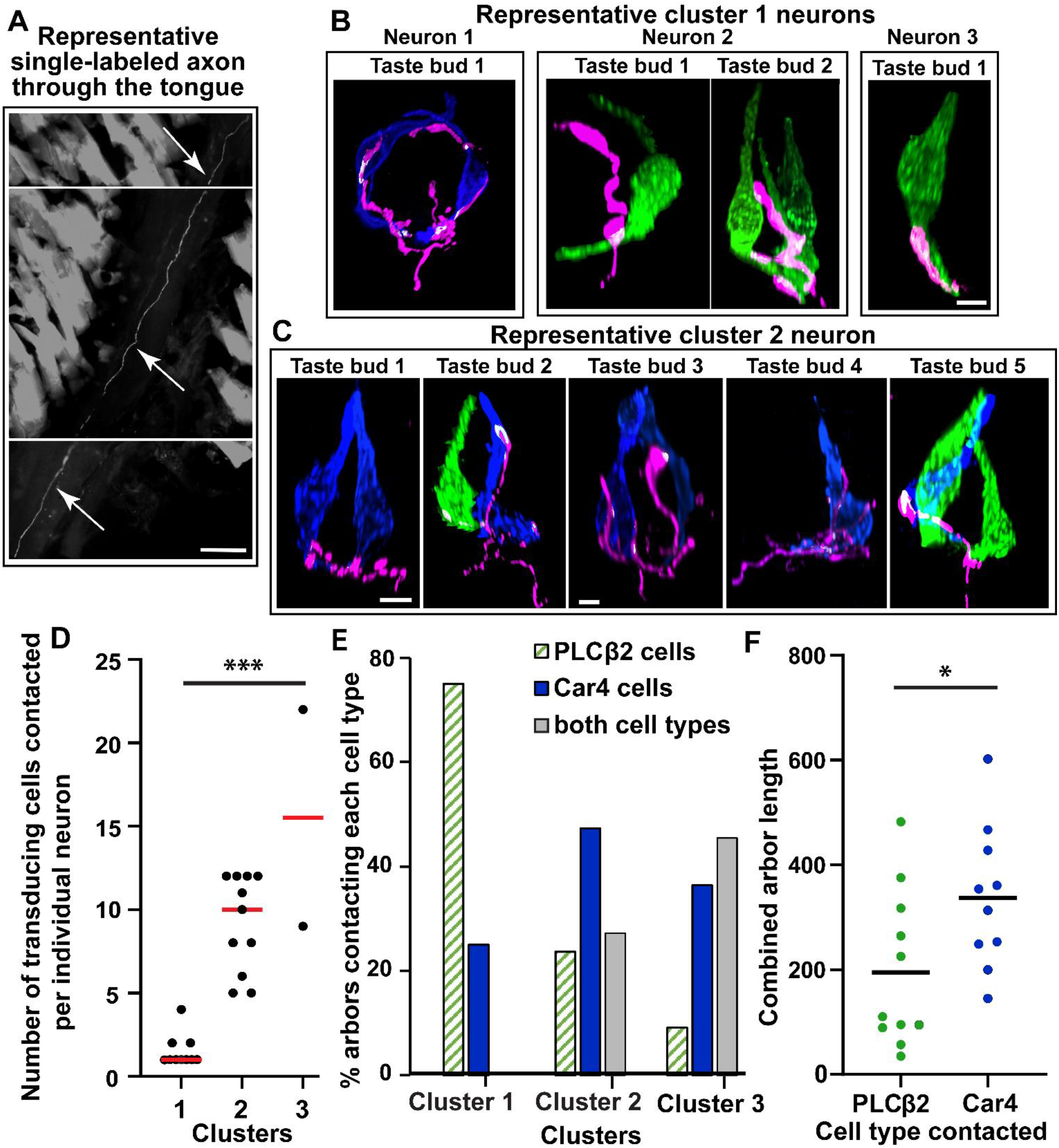
Number of taste-transducing cells contacted by axons from individual neurons. (A) A single axon (white arrows) is shown in three serial sections of tongue muscle (arrows). Scale bar = 50 µm. (B) Arbors from three cluster 1 neurons. The axon of neuron 1 (left) innervates a single taste bud with one arbor that contacts two Car4-positive cells. The axon of neuron 2 innervates two taste buds and contacts four PLCβ2-positive cells. The axon of neuron 3 contacts a single PLCβ2-positive cell. (C) One cluster 2 neuron innervating five taste buds is shown along with the 10 taste-transducing cells contacted. All scale bars = 5 µm. For all panels the arbors and labeled taste bud cells that were contacted were segmented and the fluorescence outside the reconstruction was removed, as illustrated in Fig. 8A. (D) The numbers of taste-transducing cells contacted by 21 single taste axons increases across cluster. (E) Patterns in the number of arbors contacting each cell type are different across clusters, suggesting that neuron type/s likely differ across clusters. (F) When neurons are divided into groups based of the type of cells contacted, these two groups have different but overlapping distributions of combined arbor length, suggesting that neurons contacting more Car4-positive cells (sour transducing) are also more heavily branched.

A total of 21 single axons from clusters 1, 2, and 3 were analyzed. We found that the mean number of taste-transducing cells contacted significantly differed between clusters (H_(2,20)_=17.16 *p*=0.0002; Fig. 9D). Arbors from the cluster 1 neurons are more likely to contact a PLCβ2 cell and less likely to contact a Car4 cell than arbors from clusters 2 and 3 neurons (Fig.9E, χ^2^ =6.63, p=0.036). Most cluster 1 neurons only contact one taste-transducing cell (always PLCβ2-labeled). This finding indicates that either many taste neurons are either unresponsive to stimuli or a neuron response can be driven by input from only a single taste-transducing cell. Another difference across these morphological clusters was the pattern in the type of taste-transducing cells contacted across the clusters.

We examined this same question another way using a similar approach as functional studies. Functionally, gustatory neurons are typically divided based on their “best” stimulus quality (sweet, sour, etc.), even though roughly 40% of the population responds to more than one stimulus (Frank 2000, Yoshida, Yasumatsu et al. 2006, Barretto, Gillis-Smith et al. 2015, Wu, Dvoryanchikov et al. 2015). We speculated that a similar approach would permit us to compare morphologies across predicted functional quality by dividing neurons based on the type of cell contacted most frequently, PLCβ2 (sweet/bitter/umami) or Car4 (sour) cells. For example, the neuron in Figure 9C contacted 7 Car4-positive cells and 3 PLCβ2-positive cells, so it was placed in the group contacting more Car4 cells. The two neurons that contact the same number of each cell type were placed into groups based on the combined size of these contacts. Of the 21 neurons examined, 11 contacted more sweet/bitter/umami cells, while 10 contacted more sour-transducing cells. Neurons contacting more sour-transducing cells had a greater combined arbor length (Fig. 9F; t_(19)_=2.59, p=0.018) and more terminal branch ends (median=5 (N=11) vs 16 (N=10) U=19, p=0.025), indicating that they are more heavily branched than neurons primarily contacting sweet/bitter/umami-transducing cells. However, there was also considerable variation within each group and distributions were overlapping (Fig. 9F).

## Discussion

Taste neurons exhibit both functional (Frank 1973, Lundy and Contreras 1999, Frank 2000, Sollars and Hill 2005, Yoshida, Yasumatsu et al. 2006, Breza, Nikonov et al. 2010, Yoshida and Ninomiya 2010, Barretto, Gillis-Smith et al. 2015, Wu, Dvoryanchikov et al. 2015) and molecular diversity (Yee, Yang et al. 2001, Zhang, Jin et al. 2019). Our goal was to determine if they are also morphologically diverse. Using sparse-cell genetic labeling, we found considerable variation in the branching characteristics of taste neurons. Roughly half of the taste neurons had few branches (cluster 1), whereas others branched extensively along a continuum of complexity (clusters 2-4). Some axons increased branching by innervating more taste buds (cluster 2), while others also increased the number of arbors per taste bud (clusters 3 and 4). This variability in branching complexity is surprising in mice, since it is not consistent with previous studies that utilized indirect approaches to examine the branching characteristics of taste axons (Zaidi and Whitehead 2006). This variation in branching likely underlies variation in the amount of convergence of taste-transducing cell input onto individual neurons. Consistently, we found that simple neurons (cluster 1) typically only contacted 1–4 taste-transducing cells, all of the same type. Alternatively, more heavily branched (cluster 2,3) neurons could potentially receive input from 5–22 taste-transducing cells, consistently contacting more than one type of taste-transducing cell. In addition, more heavily branched neurons were more likely to contact sour-transducing cells than those that transduce sweet/bitter/umami, indicating that neuron types (divided based on taste quality) likely differ in morphology. Lastly, variation in morphology within a neuron type or quality likely has important implications for both function and plasticity.

To what extent the variable morphological complexities represent a snap shot in time of a changing pattern or are permanent characteristics is unclear. A unique feature of taste bud cells is that they have a limited lifespan and are constantly renewed (Beidler and Smallman 1965, Farbman 1980, Delay, Kinnamon et al. 1986, Perea-Martinez, Nagai et al. 2013). As a result, gustatory ganglion neurons must continually locate and form functional connections with new adult taste-transducing cells. Presumably, this process is accompanied by changes in branching characteristics. We observed substantial variation in neuron complexity which predicts differences in the number of taste-transducing cells contacted by individual neurons. If branching changes over time, degree of convergence could also be a changing feature of taste neurons resulting in changing functional characteristics over time (Shimatani, Nikles et al. 2003). However, this may be true only within a limited range if some components of the neuron structure are stable over time.

One possibility is that arbor complexity is dictated primarily by the process of finding and connecting with a new taste-transducing cell, while arbor number is a stable feature of the neuron. We found no differences in mean terminal arbor length or complexity across clusters, consistent with the idea that arbor structure is primarily plastic. The largest/most complex arbors could be extending throughout the taste bud to locate and connect to a new taste-transducing cell. Consistently, arbors contacting multiple cell types were longer and more heavily branched. In spite of their variable morphology, most arbors only contact 1-2 taste-transducing cells, which is consistent with electron microscopy findings for synapses (Kinnamon, Sherman et al. 1988). Consistently, we found that the number of arbors is the best predictor of the number of taste-transducing cells contacted by an individual neuron. The number of arbors is determined by the number of branch points below the epithelium. Because a change in these branch points may not be required for a taste bud to form a new connection with a taste-transducing cell, the number of arbors may be a stable characteristic of the neuron. Because number of arbors determines the number of taste-transducing cells contacted this could contribute to functional stability over time.

We examined the number of taste-transducing cells contacted by each neuron. Contacts likely represent one of three scenarios, only one of which represents functional connectivity. First, there are likely some locations where the nerve arbor simply passes within 200nm of a taste-transducing cell without any specific interaction with that cell. A second possibility are locations where the cell membranes of neurons and taste-transducing cells contact as part of the process of re-innervation with cell turnover (as discussed above). Contact between taste-transducing cells and nerve arbors would allow a nerve arbor to determine molecular compatibility (i.e. the presence of a ligand on one cell and receptor on the other for factors involved in synapse formation). These “sampling contacts” are likely since the continued rewiring of the taste system is thought to depend upon multiple molecular factors (Lee, Macpherson et al. 2017). The third scenario is that all functional connections require a contact measurable at the light level. Distinguishing between these possibilities for an individual neuron is not possible even with EM analysis (Yang, Dzowo et al. 2019). This is the case because individual arbors from the same neuron, but innervating different taste buds, are separated by too great a distance to be reconstructed at the EM level. In addition, structural correlates may not exist between nerve fibers and all cell types, in spite of recent advances (Romanov, Lasher et al. 2018, Yang, Dzowo et al. 2019). However, it seems likely that the number of functional connections is greater in neurons with many arbors than for neurons with only a single arbor. If each neuron only had a functional connection on only one arbor, regardless of the number arbors, only 96 of the 452 arbors we observed would have a connection, leaving 78% of all arbors without a connection to a taste-transducing cell – a possibility which is not consistent with EM studies (Kinnamon, Sherman et al. 1988). Thus, the most likely reason for differential branching is differential convergence.

Variations in the amount of convergent input from the same type of taste-transducing cell onto different neurons could result in variable sensitivities to the taste stimulus transduced by these cells. For example, a neuron receiving input from eight sour-transducing cells (Car4 expressing (Huang, Chen et al. 2006, Chandrashekar, Yarmolinsky et al. 2009)) may be more sensitive to citric acid than a neuron receiving input from only two sour transducing cells. Consistent with this idea, stimulation of independent areas of a neuron’s receptive field with the same stimulus enhances the response (Miller 1971). Our results show that not all neurons branch to contact multiple taste-transducing cells, suggesting that this enhancement occurs in some neurons, but not others. These differences could produce variations in thresholds and intensity ranges for the same stimulus across the population of neurons. While increases in response rate represent taste stimulus intensity (Ganchrow and Erickson 1970, Scott, Plata-Salaman et al. 1991, Breza, Nikonov et al. 2010, Fonseca, de Lafuente et al. 2018), it is unclear whether additional peripheral neurons are recruited as stimulus intensity increases (i.e. graded intensity coding), as is the case with warm stimuli (Wang, Belanger et al. 2018). Variation in stimulus thresholds across the population is consistent with our anatomical data and would permit a greater range of intensities to be coded by taste neurons than by individual taste-transducing cells (Caicedo, Kim et al. 2002). This possibility is supported by the findings from the few studies that utilized multiple stimulus concentrations to examine taste coding in peripheral neurons (Ganchrow and Erickson 1970, Breza, Nikonov et al. 2010, Wu, Dvoryanchikov et al. 2015). Consistent with this idea, our data predict a range of branching characteristics within each taste quality.

We also observed that neurons contacting more sour-transducing cells tended to be more heavily branched than those contacting sweet/bitter/umami transducing cells. If as a result, neurons responding to sour stimuli receive input from a larger number of taste-transducing cells, they would be predicted to be more broadly tuned. Approximately one-fourth of individual taste-transducing cells are capable of responding to more than one stimulus (Tomchik, Berg et al. 2007, Yoshida and Ninomiya 2016). Therefore, as the amount of convergent input increases for a given neuron, the probability that it will receive input from taste-transducing cells responding to multiple stimuli also increases. Variation in branching may explain why functional studies of peripheral taste neurons have consistently observed both narrowly and broadly tuned neurons (Yoshida, Yasumatsu et al. 2006, Barretto, Gillis-Smith et al. 2015, Wu, Dvoryanchikov et al. 2015). Consistently, neurons responding to sweet are typically described as more narrowly tuned than neurons responding to primarily to sour (Lundy and Contreras 1999, Frank 2000, Breza, Nikonov et al. 2010). While it has been repeatedly speculated that this is due to differences in the tuning properties of the taste-transducing cells (Tomchik, Berg et al. 2007, Barretto, Gillis-Smith et al. 2015), our data suggest that differential convergence onto the nerve fiber is another likely explanation.

## Acknowledgements

We would like to thank Chad Samuelsen for his help with cluster analysis and comments on the manuscript. We would like to thank Kaytee Horn for her help breeding and genotyping the mice and additional technical assistance. This project was supported by R21 DC014857 and R01 DC007176 to R.F.K and F31 DC017660 to L.O.

